# 3D-Printed Magnetic Levitation Device with Deep Learning-Assisted Particle Tracking and Analysis for High-Throughput Sorting

**DOI:** 10.1101/2025.01.15.633098

**Authors:** Malavika Ramarao, Alfa Ozaltin, Sena Yaman, Kaan Ozbozduman, Irem Loc, Nil Ertok, Aria Gao, Mehmet Burcin Unlu, Naside Gozde Durmus

## Abstract

Intrinsic density-based particle separation is fundamental to biomedical research and materials science. Magnetic levitation offers an accessible and label-free approach; however, current platforms are limited by throughput, complex fabrication requirements, and manual analysis methods. Here, we demonstrate a high-throughput magnetic levitation-based microfluidic device fabricated using commercial 3D printing, integrated with dual automated analysis systems. The device features optimized magnet configurations and wide channel design (1 mm × 1 mm) that enables gentle separation (<1 PSI) at throughputs of 66 mL/hr–– a ten-fold improvement over existing levitation platforms. We developed two complementary analysis tools: “Phase” for static levitation height measurements, and a deep learning pipeline combining CNN-based particle classification (>95% accuracy) with SORT (Simple Online and Realtime Tracking) algorithm for dynamic analysis. The automated system showed excellent correlation with manual counting (Pearson coefficients: 0.91-0.99, p<0.001). Through systematic optimization of magnet spacing and paramagnetic medium concentration (150 mM Gd), the platform achieved robust continuous-flow sorting while maintaining exceptional purity (>90%) and resolving density differences as small as 0.03 g/mL. This work establishes a versatile platform for particle sorting, enabling sophisticated analysis without specialized facilities or extensive operator training, with broad applications in biomedical research and diagnostics.

## Introduction

Cell sorting has become ubiquitous across laboratories and medical centers nationwide^1–13^. Cell sorting is utilized in a wide range of procedures, from isolating specific cell populations from *in vivo* downstream analysis to capture of rare cells, such as circulating tumor cells (CTCs), from patient samples for diagnostics^14–16^. A few gold-standard methods are used for cell sorting. Examples of these methods include fluorescent-activated cell sorting (FACS) and magnetic-activated cell sorting (MACS). However, these procedures have several limitations^17,18^. For example, these techniques often require expensive equipment, extensive training, and labeling of cells with antibodies or magnetic beads respectively. Therefore, the field has started exploring alternative methods of sorting, particularly through microfluidic systems^19–23^.

Label-free sorting of particles based on their physical properties, such as size and density has been demonstrated^24^. For example, dielectrophoresis uses an electrical field, magnetophoresis employs magnetic fields, optical tweezers utilize laser beams to trap and analyze particles, and hydrophoresis leverages pressure differences to separate particles by size^25–28^. Label-free microfluidic systems usually preserve the integrity of the cell membrane and allow for accurate separation of cells with smaller sample sizes, sometimes even at the single-cell level. However, these systems still require specialized and expensive equipment that can be difficult for all labs to access and use.

In this wide range of cell sorting techniques, magnetic levitation is a promising solution to increasing accessibility for cell and particle sorting platforms. Magnetic levitation platforms can be constructed out of relatively inexpensive materials including commercially available magnets, capillaries, needles, and tubing, and these platforms can be almost immediately utilized for cell and particle sorting^27–30^. As particles traverse through a paramagnetic medium within a magnetic field, they achieve distinct levitation heights based on their intrinsic physical properties, particularly density. This enables label-free separation without compromising cellular viability or function^27–30^.

The principle of magnetic levitation relies on the balance between magnetic and gravitational forces. When introduced into a paramagnetic medium within a magnetic field, particles experience a magnetic force proportional to the difference between their magnetic susceptibility and that of the surrounding medium^31–33^. This magnetic force is coupled with both a buoyancy force dependent on the displaced volume of fluid and gravitational forces determined by the particle’s mass. This physical interaction results in density-specific equilibrium positions, enabling high-resolution separation based on subtle physical differences.

While magnetic levitation shows considerable promise, fabricating appropriate microfluidic devices has traditionally required complex manufacturing processes and specialized facilities^34,35^. Three-dimensional (3D) printing technology offers an opportunity to overcome these limitations by enabling rapid, cost-effective production of microfluidic devices. However, achieving the optical clarity necessary for precise visualization and maintaining consistent channel geometries has posed significant challenges for 3D-printed microfluidics^36–38^.

Here, we present a 3D-printed microfluidic device specifically designed for continuous magnetic levitation-based particle separation. By employing a FormLabs 3+ printer with Clear Resin V4 and developing optimized post-processing protocols, we demonstrate the feasibility of producing optically clear devices suitable for precise particle tracking and separation. We validate our system using polymer beads of varying densities (1.02, 1.06, and 1.09 g/mL) and utilize a custom Python-based tracking algorithm for automated analysis of separation efficiency.

This work addresses several key challenges in the field: first, it establishes 3D printing as a viable fabrication method for magnetic levitation devices, significantly reducing barriers to entry for laboratories interested in implementing this technology. Second, it demonstrates the integration of automated particle tracking and analysis, enhancing the quantitative rigor of separation performance assessment. Finally, it provides a thoroughly validated platform that can be readily adapted for various biomedical applications, from basic research to diagnostic applications in resource-limited settings.

## Materials and Methods

### 3D-Printed Magnetic Levitation Device Fabrication and Assembly

The magnetic levitation devices were fabricated using a FormLabs 3+ printer with Clear Resin V4 (FormLabs, Form 3). After printing, a systematic post-processing protocol was implemented to achieve optimal optical clarity and structural integrity. The 3D-printed levitation devices underwent sequential isopropyl alcohol baths: first for 15 minutes with active air flushing to remove excess resin, followed by a 5-minute secondary bath for final cleaning. Post-cleaning, the devices were cured at 60°C for 3 minutes and any warping was fixed using the SMD rework gun at 170°F.

A multi-step surface finishing process was developed to achieve the optical clarity required for precise particle tracking. Initial surface smoothing was performed using 800-grit sandpaper (MAXMAN) to eliminate major surface irregularities, followed by fine polishing with 1000-grit sandpaper (AutKerige). Upon completing the sanding process, the device was washed with DI water to remove any sanding residue and was allowed to air dry. After drying, a light application of Clear Acrylic Coating Spray (Krylon, 1303) was sprayed onto the device’s surface for a polished finish.

### High-Throughput Sorting Device Fabrication and Assembly

The fluidic network was established using a dual-diameter tubing configuration to optimize flow stability and enable precise flow control. Four sections of Tygon 2375 tubing (outer diameter: 0.125 inches, inner diameter: 0.0625 inches) were cut to a minimum length of 3 inches and inserted directly into the device’s inlet and outlet ports to serve as primary flow channels. These primary channels were then interfaced with smaller diameter Tygon Microbore tubing (outer diameter: 0.06 inches, inner diameter: 0.02 inches) to facilitate connection with external flow control systems and enable precise sample handling. All tubing connections were permanently secured using medical-grade epoxy (Devcon, 14250) to ensure system integrity and prevent leakage during high-throughput sorting. This microfluidic design maintains stable flow dynamics throughout the device while operating at flow rates up to 66 mL/hr, with an approximate internal volume of 150 μL per channel. The design of this fluidic network enables consistent and reliable particle sorting performance while minimizing dead volume and sample loss.

### Simulation of the Magnetic Field and Particle Levitation

Simulations of the magnetic field and particle levitation height were conducted using the Magpylib package (Python version 3.10.12) and a finite element modeling (FEM) tool (COMSOL version 5.5). The magnetic induction generated by two N52-grade NdFeB magnets was modeled in 2D with a magnetization of 1150 kA/m. Magnetic flux density profiles were analyzed for various device designs.

Due to the inherent variability in 3D printing and fitting the permanent magnets into the devices, there are differences in the distance between the magnets from the design to the printed device. The distance between the magnets was measured with calipers (Neiko, 01407A), and this exact parameter was used for all simulations. The average distance between the magnets were measured as 3.180 mm, 3.495 mm, and 3.765 mm for Device 1, Device 2, and Device 3 respectively. For all data analysis and simulations concerning the static levitation height of the polyethylene beads, these experimental values are used in order to obtain accurate results.

To simulate particle levitation, the fluid density was adjusted to reflect the addition of gadolinium (Gd), with densities set to 1.039 g/mL, 1.054 g/mL, and 1.068 g/mL for Gd concentrations of 100 mM, 150 mM, and 200 mM, respectively. In the simulations, the magnetic susceptibility of Gd and polyethylene beads were taken as 3.2 x 10^-4^ M^-1^ and 0, respectively. The simulated levitation heights of particles with densities of 1.02, 1.06, and 1.09 g/mL were reported as their distances from the top surface of the bottom magnet.

### Static Levitation and Quantification of Levitation Heights

Static levitation experiments were conducted using density-calibrated polymer microspheres (Cospheric) of three distinct densities: 1.02 g/mL (180507-110-1), 1.06 g/mL (180507-110-3), and 1.09 g/mL (180507-110-5). Sample preparation consisted of mixing a 1 mL solution generated by suspending the beads in PBS containing 0.1% Pluronic and varying concentrations of Gadavist (McKesson, 1194924) as the paramagnetic agent. Samples were then pulled into the capillary at a constant flow rate using 10 mL syringes (BD, 302995) connected via 24-gauge needles (Fisnar, 8001127) to the outlets and controlled via syringe pumps (InfusionONE, 98242). Levitation heights were monitored by manually taking images using standardized camera settings at approximately 30-second intervals, and images were quantified using our custom Phase analysis program, as found on github.com/vxco/phase.

### High-Throughput Flow Sorting and Quantification of Sorting Efficiency

For continuous sorting experiments, 1.02 g/mL, 1.06 g/mL, and 1.09 g/mL beads (Cospheric, DMB-FGRN-1.02, DMB-FVIO-1.06, and DMB-FRED-1.09 respectively) were mixed in a solution of PBS with 0.1% Pluronic and 150 mM Gadavist (Gd). The device was equilibrated with a 150 mM Gadavist solution before the sample introduction. The continuous flow was maintained using three syringe pumps (InfusionONE, 98242) connected to the outlet ports.

To optimize the sorting conditions, initial experiments were performed in each device, testing different flow rates while observing sorting accuracy in real time. Based on these initial tests, the flow rates were gradually adjusted to determine the ideal sorting conditions within each device. In Device 1 the flow rates were set at 500 ul/min, 100 ul/min, and 400 ul/min at the top, middle and bottom outlets, respectively. In Device 2, the flow rates were set at 500 ul/min, 170 ul/min, and 400 ul/min at the top, middle, and bottom outlets, respectively. In Device 3, the flow rates were set at 500 ul/min, 200 ul/min and 400 ul/min at the top, middle, and bottom outlets, respectively.

The mixed population sample containing the three bead types was introduced into the device, where the beads were sorted into three outlets based on their densities. During each trial, 3-4 mL of the mixed population sample was processed. When approximately 90% of the mixed population sample had been sorted, an additional 3-4 mL of 150 mM Gadavist solution was gradually added to the sample to ensure no bubbles formed within the tubing before the bead sorting procedure was fully completed. The entire sorting process was documented using an Apple iPhone 14 (4K resolution at 60 fps) positioned 10 cm from the device, capturing a 4.8 mm x 2.7 mm field of view under controlled LED back-lighting (1000 lux). This standardized video acquisition protocol enabled both automated particle tracking analysis and manual verification of sorting efficiency. For manual quantification of sorting efficiency, sorted fractions were collected in a standard 6-well plate and centrifuged at 2,000g for 3 minutes to concentrate the beads. Wells were imaged against both white and black backgrounds to enhance particle contrast. Particle counts were performed using ImageJ Multi-Point tool, with sorting purity and recovery rates calculated as:

Purity = (Correct particles in the outlet / Total particles in the outlet) × 100%

Recovery = (Correct particles in the outlet / Total input particles of that type) × 100%

### Bead Sorting Tracking Analysis

Video frames were processed using a custom Python pipeline developed with OpenCV to perform background subtraction, contour detection, and color classification. A Mixture of Gaussians background subtraction algorithm was applied to isolate moving beads from the static background. This method generated a foreground mask by identifying pixel intensity changes across frames. The contours of the segmented beads were extracted from the foreground mask and used to crop regions of interest (ROIs) corresponding to individual beads.

To classify bead colors, the cropped ROIs were resized to 128×128 pixels and passed to a convolutional neural network (CNN) model trained on 5,000 labeled images for each of the four categories: 1.02 g/mL (green), 1.06 g/mL (purple), and 1.09 g/mL (red) “beads” and “no bead”. The “no bead” category accounted for noise and artifacts that did not correspond to the valid beads such as small dust particles, and such detections were discarded from the processing pipeline. The CNN, designed with three convolutional layers and two fully connected layers, analyzed the spatial and color features of the ROIs and assigned each bead to its corresponding category. This approach offered improved accuracy and robustness to variations in lighting and background conditions.

The SORT (Simple Online and Realtime Tracking) algorithm was employed for consistent object identification across frames^39^. SORT uses a Kalman filter to predict bead positions and the Hungarian algorithm to associate detections with existing tracks based on bounding box Intersection over Union (IoU). Each bead was assigned a unique ID by the SORT tracker, enabling the system to maintain consistent labeling across frames. This approach ensured robust and efficient tracking of individual magnetic beads over time.

The final analysis was conducted on the accumulated data from SORT tracking and CNN-based classification to evaluate the bead sorting efficiency of the magnetic levitation sorting device. Beads were classified into three distinct outlets (top, middle, and bottom) based on their tracked positions within the device. For each outlet, the total bead counts were determined for each color category (green, purple, red). Outlet purity and recovery were subsequently calculated. The annotated frames were saved to an output video file, providing real-time visual feedback on bead trajectories and classifications.

## Results and Discussion

### 3D-Printed Magnetic Levitation Device Design and Development

Current gold-standard sorting devices such as FACS and MACS face significant limitations in terms of cost, complexity, and infrastructure requirements. While emerging technologies like dielectrophoresis and optical tweezers offer promising alternatives, they still require expensive equipment and cleanroom facilities for fabrication. To address these challenges in the field, we have developed a cost-effective 3D-printed magnetic levitation platform that enables high-throughput, gentle particle sorting, based on density (**Figure 1**). This innovation in microfluidics and sorting technology will expand access to sorting technology, overcoming barriers of entry in the field.

**Figure 1.**
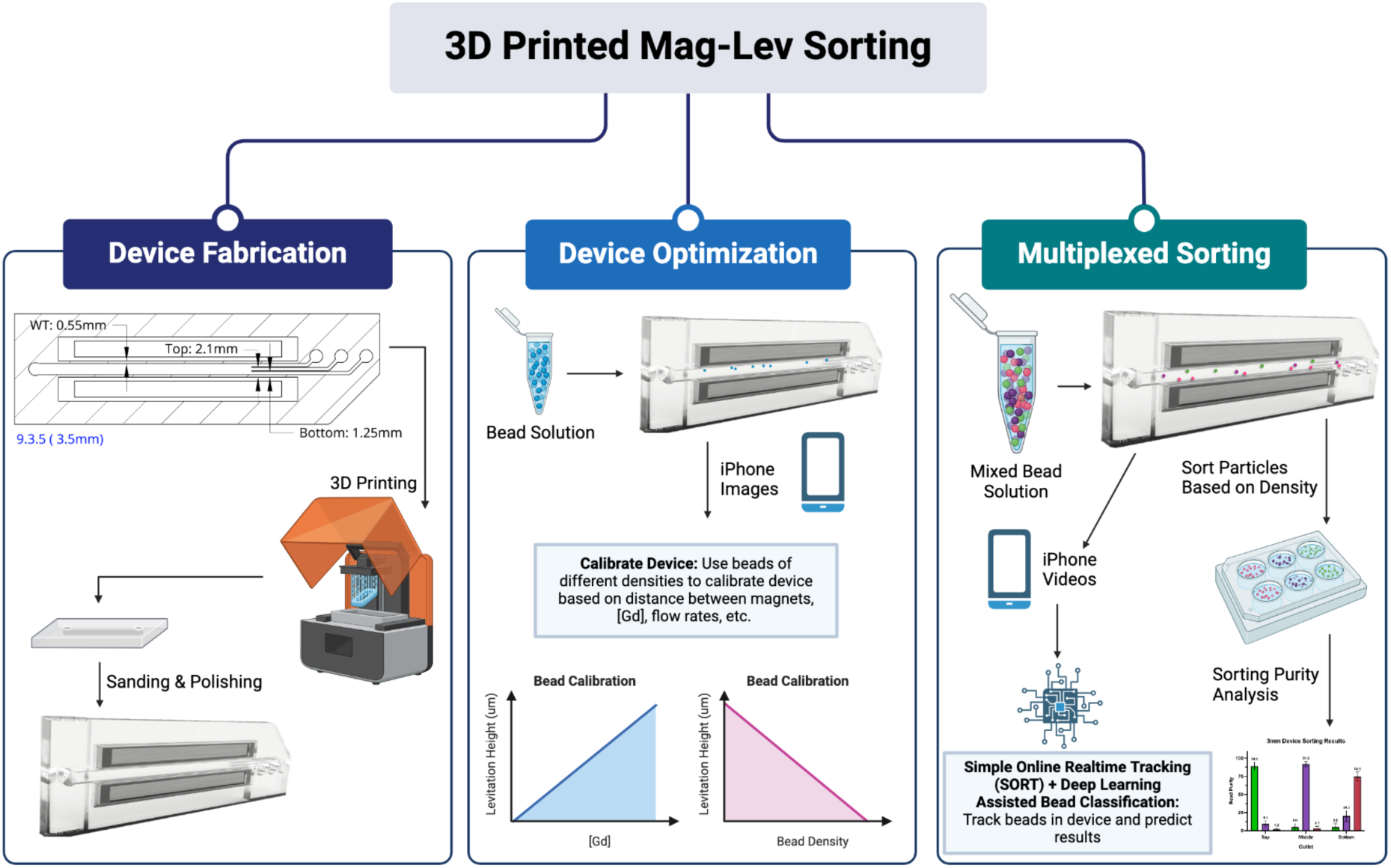
A 3D-printed Magnetic Levitation Platform with Automated Particle Tracking for Multiplexed Sorting. We demonstrate a high-throughput magnetic levitation-based microfluidic device fabricated using commercial 3D printing, integrated with dual automated analysis systems. (Created with Biorender.com).

The 3D-printed magnetic levitation device consists of the printed frame with inbuilt capillary and magnet holders, two permanent neodymium bar magnets (50 mm length, 2 mm width, and 5 mm height), and Tygon tubing. The permanent magnets fit securely into the magnet holders built into the device frame, and the tubing connects the inlets and outlets of the device to the sample and syringe pumps respectively for real-time sorting operations (**Figure 1**). Additionally, to accommodate real-time imaging of static levitation and sorting of particles, a frame was built to hold the 3D printed device and a phone so images and videos of particles within the device could be procured.

Using parametric CAD design, we investigated three device designs with different inter-magnet spacings (3.0, 3.25, and 3.5 mm) to optimize field strength and sorting performance. This modular approach enabled rapid design iteration while demonstrating the platform’s adaptability. Using a 3D printer from Formlabs, the devices were designed to minimize the distance between the permanent magnets to maximize the magnetic field experienced by the particles within the chamber. For alternate sorting applications, the dimensions and magnetic field can be easily altered through adjustments in the device design. Overall, this new magnetic levitation design integrates several key engineering innovations: a) a wider microfluidic channel (>1 mm × 1 mm cross-section) designed to minimize physical stress on particles, operating at <1 PSI compared to 20-30 PSI in conventional FACS systems, b) an extended flow path (5 cm) optimized for enhanced separation resolution, c) precisely engineered magnet holders maintaining optimal field geometry, and d) custom-designed imaging mount compatible with standard smartphones enabling real-time particle tracking and monitoring (**Figure 1**).

### Simulated Levitation of Particles in the 3D-Printed Magnetic Levitation Device

Magnetic levitation is a robust and unique method for sorting particles, including cells, based on their density and biophysical properties. When particles are mixed in a paramagnetic medium (in this case, gadobutrol, a non-ionic, FDA-approved paramagnetic solution containing Gd3+) and placed within a magnetic field, they will levitate to a unique equilibrium height based on their density. At this equilibrium levitation height, the gravitational force (Fg) is equal and opposite to the buoyancy force (Fb) and the magnetic induction force (Fm) acting on the particles. We systematically investigated the effect of magnetic field geometry on separation performance using three device configurations with permanent magnet spacings designed to be 3.0 mm (Device 1), 3.25 mm (Device 2), and 3.5 mm (Device 3).

To optimize device performance, we conducted comprehensive COMSOL and Magpylib simulations of magnetic field distributions within the channel (**Figure 2A, 2B**). To evaluate the capability of our 3D-printed magnetic levitation devices we used 1.02 g/mL, 1.06 g/mL, and 1.09 g/mL polyethylene beads as model particles. We simulated levitation heights and separation distances across different paramagnetic concentrations using COMSOL (**Figure 2C, 2D**).

**Figure 2.**
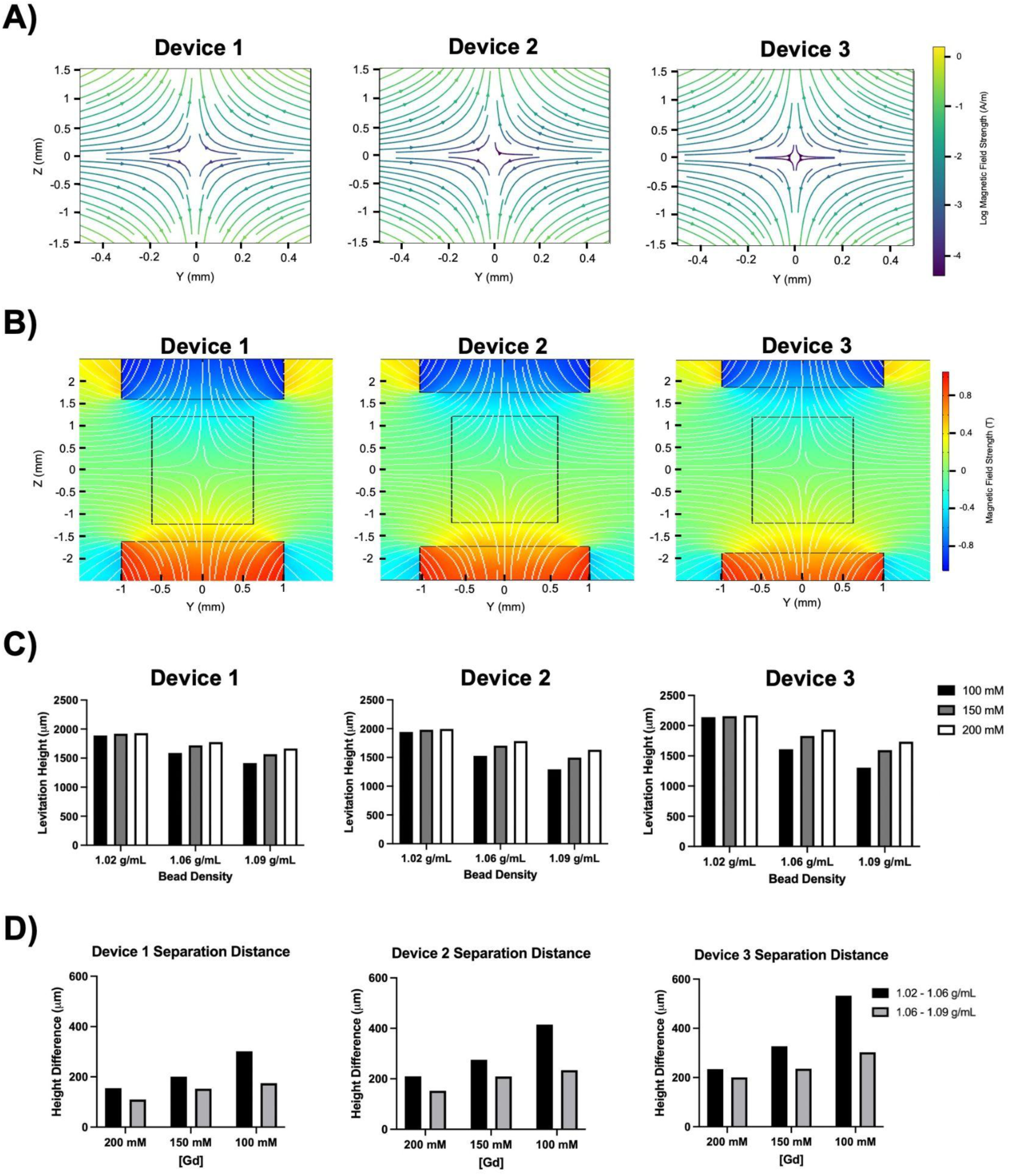
Simulated Magnetic Field and Levitation Height of Particles. **A)** Magnetic field lines within the capillary for devices with different distances between the magnets. **B)** Simulated magnetic field in the device for devices with different distances between the magnets. **C)** Simulated levitation height of beads of differing densities within magnetic levitation devices at different Gadavist concentrations. **D)** Simulated separation distance between beads of differing densities within magnetic levitation devices at different Gadavist concentrations.

The simulated levitation height of the beads at 150 mM Gd were 1919.86 μm, 1719.23 μm, and 1566.67 μm in Device 1, 1980.53 μm, 1705.03 μm, and 1496.00 μm in Device 2, and 2157.68 μm, 1830.26 μm, and 1594.65 μm in Device 3 for the 1.02, 1.06, and 1.09 g/mL polyethylene beads respectively (**Figure 2C**). These levitation heights are calculated as the distance from the bottom permanent magnet, and the height differs based on both the magnetic field applied to the particles and the paramagnetic medium concentration. The simulated separation distance between the beads was also calculated to predict the best conditions for sorting the beads (**Supplementary Figure 1**). The separation distance between the 1.02 g/mL and 1.06 g/mL beads at 150 mM Gd were 200.63 μm, 275.50 μm, and 327.42 μm for Devices 1, 2, and 3 respectively (**Figure 2D**). The separation distance between the 1.06 g/mL and 1.09 g/mL beads at 150 mM Gd were 152.56 μm, 209.03 μm, and 235.61 μm for Devices 1, 2, and 3 respectively (**Figure 2D**). The separation distance of the different density beads is affected by the Gd concentration and the magnetic field applied to the particles. These values were used to determine the best conditions for static levitation and sorting of the polyethylene beads.

### Static Levitation Characterization

To validate simulation results, we developed “Phase”––a custom image analysis software for automated tracking of particle levitation heights. The software utilizes the permanent magnets as fiducial markers to calibrate imaging parameters and enables precise measurement of particle positions within the channel (**Figure 3A, 3B**). As beads were passed through the devices at various Gd concentrations, images were taken from a standard phone camera (**Figure 3A**). The Phase program calibrated the zoom and angle of the camera using the permanent magnets as stable guides; the program user then clicked the individual beads to determine their levitation height within the channel (**Figure 3B**).

**Figure 3.**
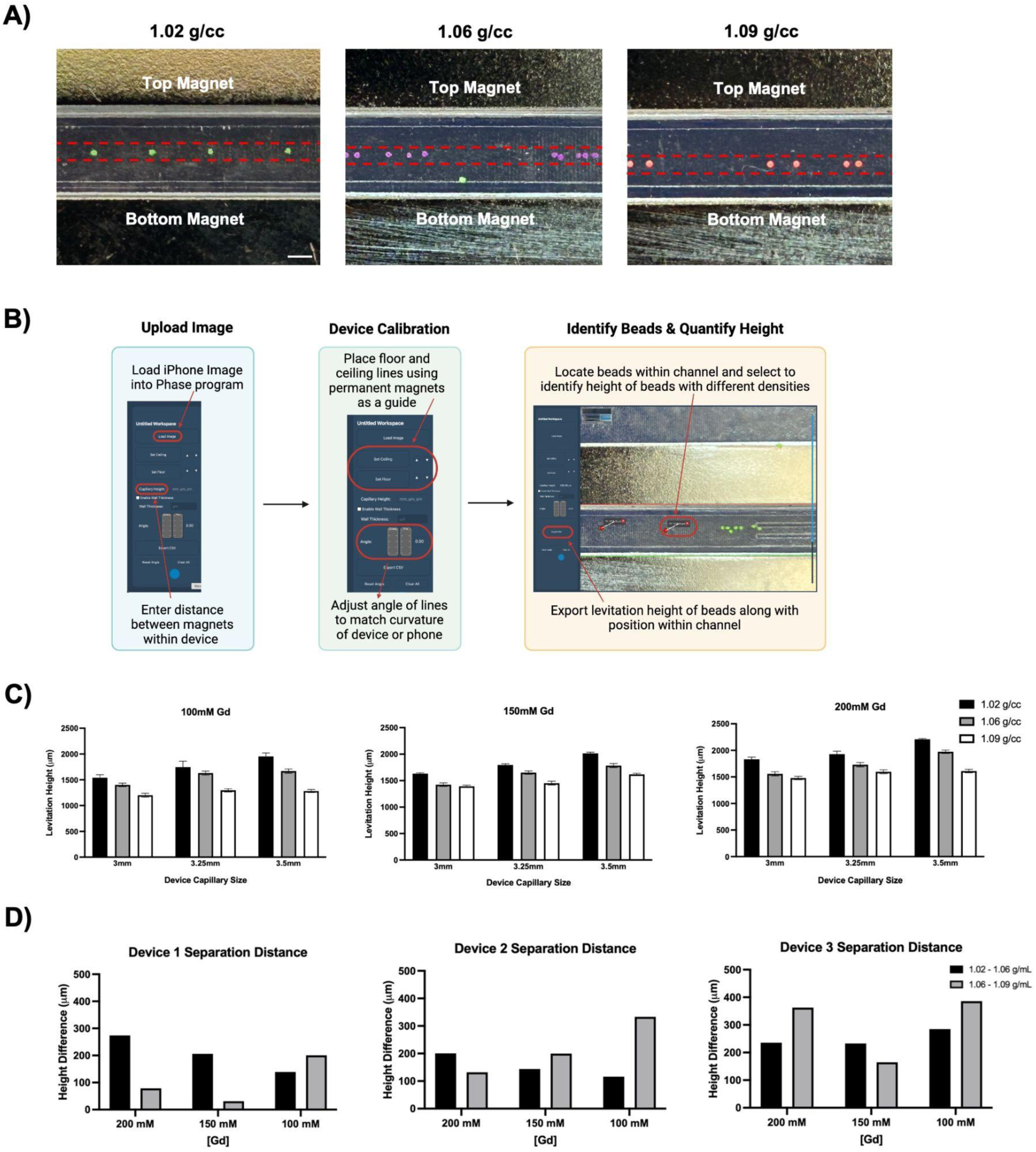
PHASE Program for Automated Levitation Height Analysis. **A)** Images of beads within the capillary of the 3.25 mm mag-lev 3D printed device at 150 mM Gd used for static levitation analysis. Scale bar is 1 mm. **B)** PHASE program data analysis pipeline used to characterize the static levitation heights of beads through capillary alignment and bead selection. **C)** Static levitation heights of beads in the device with differing Gadavist concentration and distance between the permanent magnets. Error bars represent the standard deviation of the mean. **D)** Separation distance between beads of differing densities within magnetic levitation devices at different Gadavist concentrations.

After quantifying the levitation height of twenty beads from a minimum of 4 photos per condition, the levitation profiles of the 1.02 g/mL, 1.06 g/mL, and 1.09 g/mL polyethylene beads within the 3D printed magnetic levitation devices were determined. The levitation height of the beads at 150 mM Gd were 1630.19 ± 18.29 μm, 1424.00 ± 33.87 μm, and 1392.87 ± 20.80 μm in Device 1, 1795.87 ± 22.45 μm, 1651.72 ± 31.88 μm, and 1452.01 ± 37.38 μm in Device 2, and 2015.78 ± 20.38 μm, 1783.26 ± 39.14 μm, and 1618.27 ± 20.69 μm in Device 3 for the 1.02, 1.06, and 1.09 g/mL polyethylene beads respectively (**Figure 3C**). For every bead, the levitation height was significantly correlated with both the device used and the Gd concentration (two-way ANOVA for each bead, p <0.001 for all row and column factors).

After determining how to adjust the levitation height of the polyethylene beads by changing key parameters including the magnetic field and the paramagnetic medium concentration, the ideal sorting conditions were predicted by calculating the experimental separation distance between the beads with varying parameters (**Figure 3D**). Using the static levitation heights of the beads as a guide, the 100 mM Gd and 200 mM Gd concentrations were characterized as non-ideal for the sorting of the beads. Despite the increased separation distance between the beads at these Gd concentrations, the beads were positioned too low at 100 mM Gd and too high at 200 mM Gd in the capillary to enter the correct outlets. At 150 mM Gd, the separation distance between the 1.02 and 1.06 g/mL beads as well as the 1.06 and 1.09 g/mL beads was 206.19 μm and 31.13 μm in Device 1, 144.15 μm and 199.71 μm in Device 2, and 232.52 μm and 165.00 μm in Device 3 (**Figure 3D**). Even at lower magnetic field strengths, high separation between beads of different densities was achieved. For instance, the system achieved separation distances of up to 232.52 μm between adjacent particle populations sufficient for high-purity sorting while maintaining gentle operating conditions. Thus, using the static levitation images and the Phase analysis software, the ideal sorting conditions could easily be determined using a minimal amount of sample.

### High-Throughput Multiplexed Sorting

Next, we implemented continuous-flow sorting using optimized conditions (150 mM Gd, device-specific flow rates) based on static levitation results. 1.02, 1.06, and 1.09 g/mL beads were sorted into the three outlets with high accuracy, and videos were taken of each sorting procedure using a standard phone camera (**Figure 4A, Supplementary Video 1**). The flow rates in each device were optimized separately to determine the ideal flow rates and Gd concentration for each device. Based on the flow rates utilized in each device, samples could be processed at a rate of 60 mL/hour (Device 1) to 66 mL/hour (Device 3). After determining the ideal sorting conditions and performing the multiplexed sorting, the beads were pushed out of each outlet onto a 6-well plate and manually counted using ImageJ to quantify the sorting efficacy in terms of both purity and recovery rate (**Figure 4B, Supplementary Figure 2**). The starting sample had an average distribution of 24.85 ± 10.23% 1.02 g/mL beads, 48.44 ± 9.23% 1.06 g/mL beads, and 26.71 ± 8.84% 1.09 g/mL beads across all trials. After sorting with Device 1, ending purities of 88.93 ± 5.54% 1.02 g/mL beads, 91.88 ± 3.57% 1.06 g/mL beads, and 74.5 ± 6.29% 1.09 g/mL beads were achieved (n = 4). After sorting with Device 2, ending purities of 93.91 ± 2.85% 1.02 g/mL beads, 94.40 ± 3.55% 1.06 g/mL beads, and 78.10 ± 10.01% 1.09 g/mL beads were achieved (n = 4). After sorting with Device 3, ending purities of 94.07 ± 5.52% 1.02 g/mL beads, 96.76 ± 1.78% 1.06 g/mL beads, and 77.22 ± 13.78% 1.09 g/mL beads were achieved (n = 8) (**Figure 4C**).

**Figure 4.**
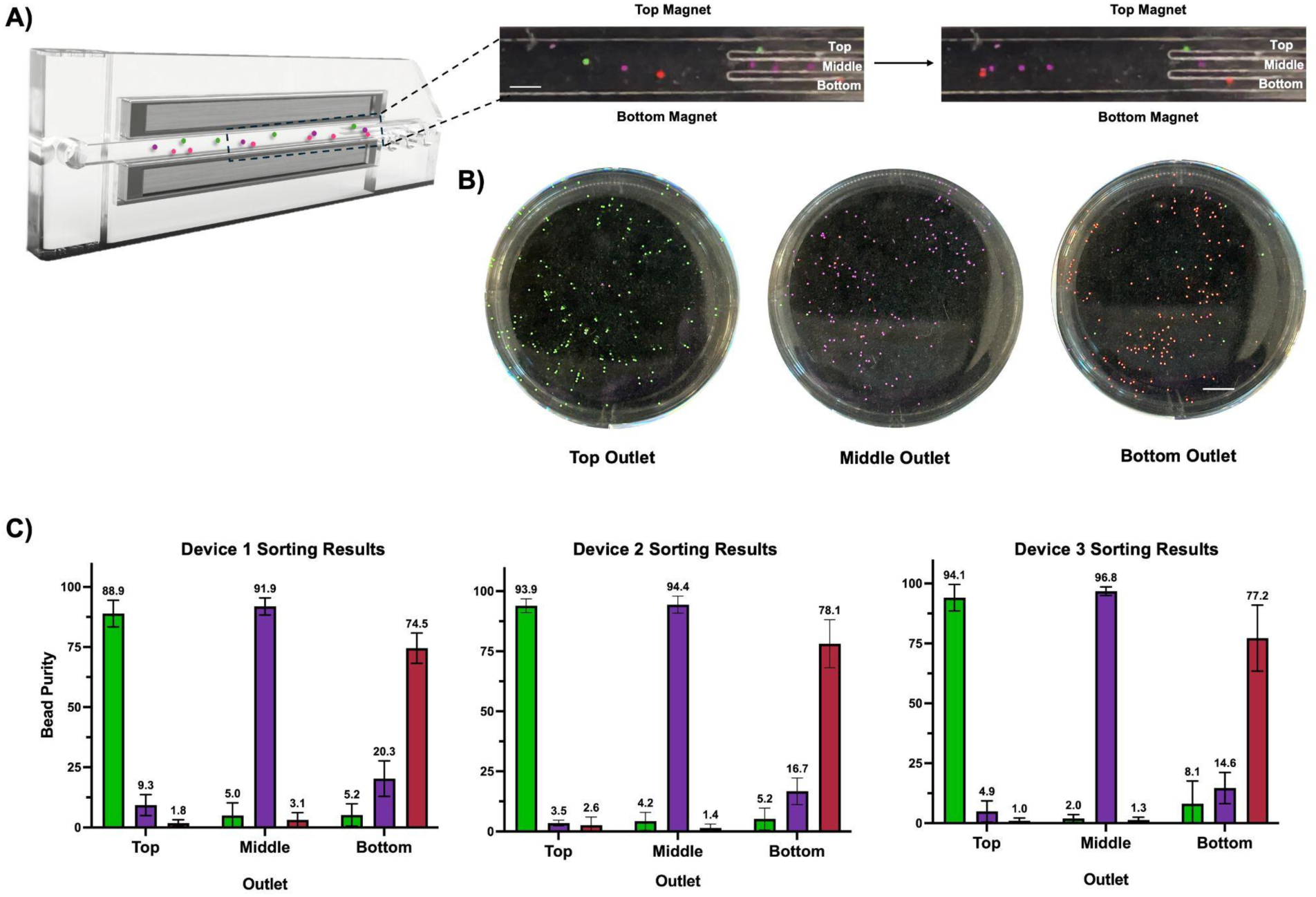
3D-Printed Mag-Lev Device Particle Sorting Results. **A)** Experimental setup for 3D-Printed Mag-Lev sorting with a zoomed-in view of particles passing through the capillary during sorting. Images were taken from Device 3, tracking trial 1 video, with the right image showing the capillary one second after the left image. Permanent magnet and outlet positions are labeled. Scale bar is 1 mm. **B)** Visualization of particles collected from each outlet and ejected onto a 6-well plate following sorting. These images depict the results from the Device 3, tracking trial 5. Scale bar is 5 mm. **C)** Purity results for each outlet following sorting at 150mM Gd and optimized flow rates for the Device 1 (n=4), Device 2 (n=4), and Device 3 (n=8). Error bars represent the standard deviation of the mean.

Thus, as predicted by the simulation data, high purity after sorting was achieved using the 150 mM Gd concentration with varying flow rates within each device. These sorting results demonstrate the ability of the 3D-printed magnetic levitation device to sort three different particle types based on density with high accuracy. Additionally, given the high flow rates used for sorting, a single printed device can process 66 mL/hr of particle solutions, which makes this device particularly useful for high-throughput high volume sorting. This represents a significant advance over existing microfluidic magnetic levitation sorters which typically process <10 mL/hour ^27–30^.

### Real-Time Tracking of Levitated Particles and Automated Analysis of Sorting Efficiency

The analyzed video files were approximately 5 minutes long, with a resolution of 1,400 x 150 pixels after cropping to focus on the flow channel and outlets. The SORT tracker was employed with an Intersection over Union (IOU) threshold of 0.1 to ensure optimal matching of bead detection across frames. (**Figure 5A**) For the underlying mixture of Gaussians-based background subtraction algorithm, a background history of 50 to 100 frames was optimized, depending on the flow rate of the magnetic levitation device. The essential attributes of the tracked ROIs are systematically logged for each video frame including the spatial coordinates of the ROI’s bounding box, size, and velocity, computed across the adjacent frames.

**Figure 5.**
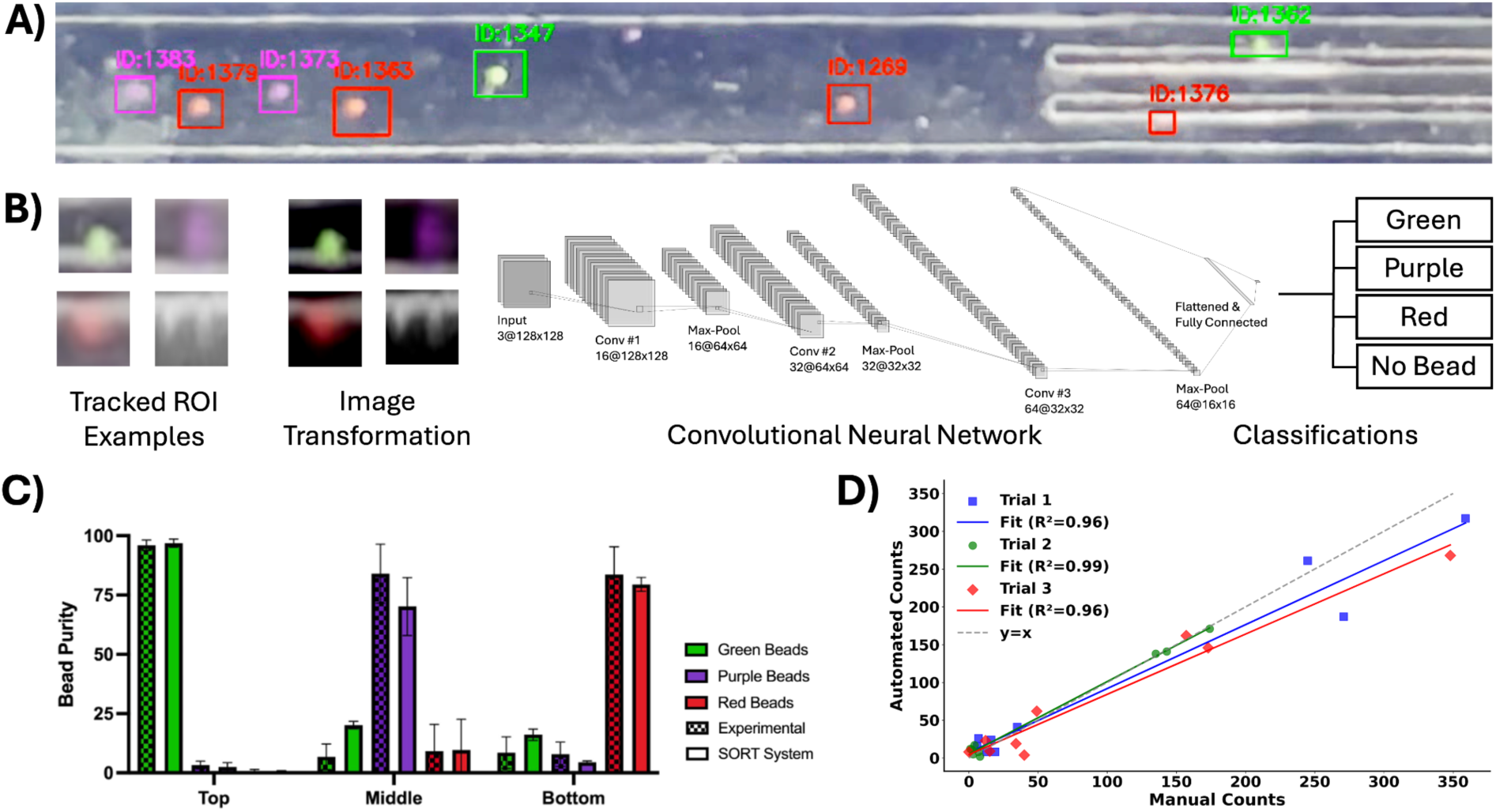
Real-time particle tracking, classification, and automated analysis of sorting efficiency. **A)** An example frame showing bead tracking results with SORT tracker across the device flow channel and outlets. Unique IDs are assigned by SORT, while bounding box colors —green, purple, or red—indicate the automated bead classifications. **B)** Preprocessing pipeline of the tracked regions of interest (ROIs) through resizing and color normalization, followed by the convolutional neural network (CNN) architecture used for bead classification into green, purple, red, or “no bead” categories. **C)** Comparison of experimental bead purity across outlets (top, middle, bottom) using manual ground truth versus automated results for each bead category. **D)** Scatter plot of manual versus automated bead counts for three experimental trials, with linear regression fits and high correlation coefficients (R² > 0.96), demonstrating strong agreement between methods.

The logged bounding boxes undergo a preprocessing pipeline before passing through the convolutional neural network (CNN). Each bounding box is cropped from the frame, resized to 128 x 128 pixels, and preprocessed through a color normalization pipeline, resulting in pixel values scaled to the range [-1, 1] (**Figure 5B**). This preprocessing standardizes the input data, allowing the CNN to focus on learning the distinguishing features of each bead category.

The CNN model features three convolutional layers with ReLU activations, followed by max-pooling for efficient feature extraction. Fully connected layers process these features, and the softmax output classifies the bead color categories while effectively filtering out noise and small dust particles (“No bead” category) (**Figure 5B**). All evaluated metrics, including accuracy, precision, recall, and F1-score, consistently exceeded 95% during the model validation, underscoring the model’s robustness and reliability. Notably, this high performance was achieved under challenging imaging conditions characterized by vibrations, the microscopic scale of the regions of interest, and the use of a commercial smartphone camera without professional optical enhancements, underscoring the effectiveness of leveraging deep learning-based methodologies to provide accurate and reliable classifications even in constrained and non-ideal imaging environments.

The valid beads exiting through each outlet were counted at multiple checkpoints along the outlet. To achieve a more precise estimation of the actual counts, the maximum value of these counts was utilized. When compared to the manual counts, which served as the ground truth, the automated system showed a high correlation with Pearson correlation coefficients of 0.978 for Trial 4, 0.997 for Trial 5, and 0.979 for Trial 6 (all with p < 0.001) (**Figure 5C, 5D**). The outlet purity rates were also highly correlated among the automated and manual counting pipelines with a correlation coefficient of 0.910 and p < 0.001. Statistical analysis showed no significant difference between the automated and manual counting methods, further validating the effectiveness of the automated system.

## Conclusion

In summary, we have successfully developed and validated a high-throughput 3D-printed magnetic levitation platform that fundamentally transforms particle separation technology. By combining commercial 3D printing with optimized magnetic levitation design, we achieved robust particle separation in a simplified, single-step manufactured device that eliminates the need for cleanroom facilities. The platform demonstrates exceptional performance, processing samples at up to 66 mL/hr while maintaining >90% sorting purity and precisely separating particles with density differences as small as 0.03 g/mL. This performance is possible due to the robust materials used for device construction, the increased capillary size, and the increased number of outlets. This sample processing rate is a marked improvement over current magnetic levitation microfluidic devices that contain only two outlets and typically use flow rates of 20-100 uL/min, resulting in a total sample processing rate of approximately 6-10 mL/hr ^27–30^. We complemented this high-throughput operation with advanced analysis capabilities through our custom Phase application for levitation height tracking and automated particle tracking, enabling real-time sorting analysis and monitoring of multiple particle parameters across different magnetic field configurations. The integration of wide channels, optimized field geometry, and automated tracking enables our platform to achieve both high-throughput and high-purity separation while maintaining exceptionally gentle sorting conditions––critical for preserving particle integrity. The system’s ability to process large sample volumes (>60 mL/hour) at low pressure (<1 PSI) while maintaining separation resolution makes it particularly suitable for applications requiring both speed and precision.

The practical advantages of our approach significantly advance the accessibility of particle sorting technology. By dramatically reducing device costs and eliminating the need for specialized facilities or extensive operator training, we have created a truly democratized platform. The use of 3D printing enables both rapid prototyping for design iteration and straightforward scalability through parallel operation. Our system’s combination of high-throughput capability, label-free operation, and cost-effectiveness opens new possibilities for researchers to implement multiplexed sorting protocols that were previously limited by technical and financial barriers.

This platform provides a foundation for transformative developments in biomedical applications. Future work will focus on optimizing additional 3D-printed devices specifically for biological materials, expanding from basic cell sorting to sophisticated clinical diagnostics. The integration of automated analysis tools and smartphone-compatible imaging makes this technology particularly valuable in resource-limited settings where traditional sorting equipment is inaccessible. By successfully combining high performance with unprecedented accessibility, our work establishes a new paradigm for particle separation technology and lays the groundwork for continued innovation in biomedical research and sample processing applications. This achievement represents not just a technical advance, but a significant step toward making sophisticated particle analysis available to a broader scientific community.

## Data Availability

The data supporting this article has been included as part of the Supporting Information. The code for the PHASe program can be found at https://github.com/vxco/phase, and PHASe version 4.0.1 was employed for this study.

## Supplementary Information

**Supplementary Figure 1.**
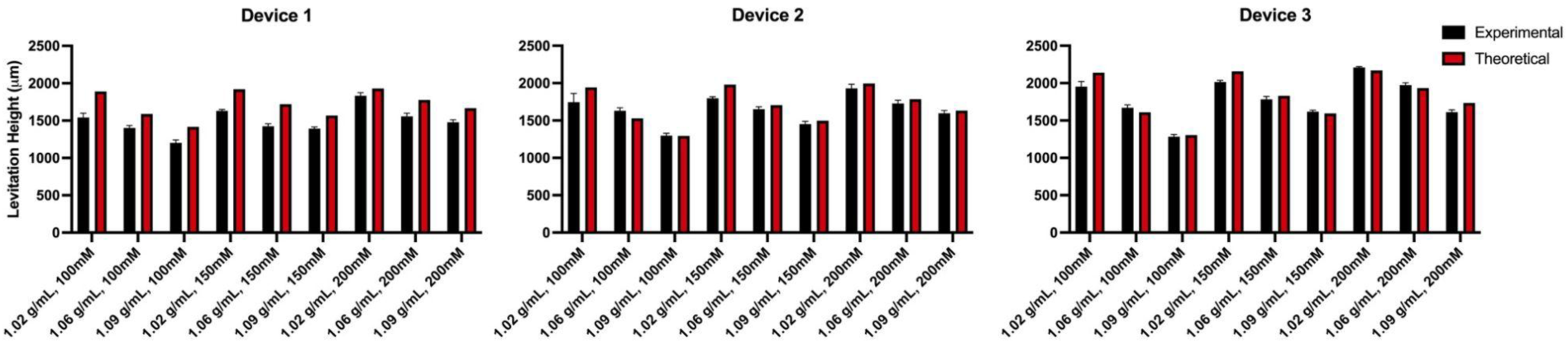
Experimental versus Simulation Static Levitation Heights. Comparison of simulated versus experimental static levitation heights of different density beads at varying Gadavist concentrations and devices.

**Supplementary Figure 2.**
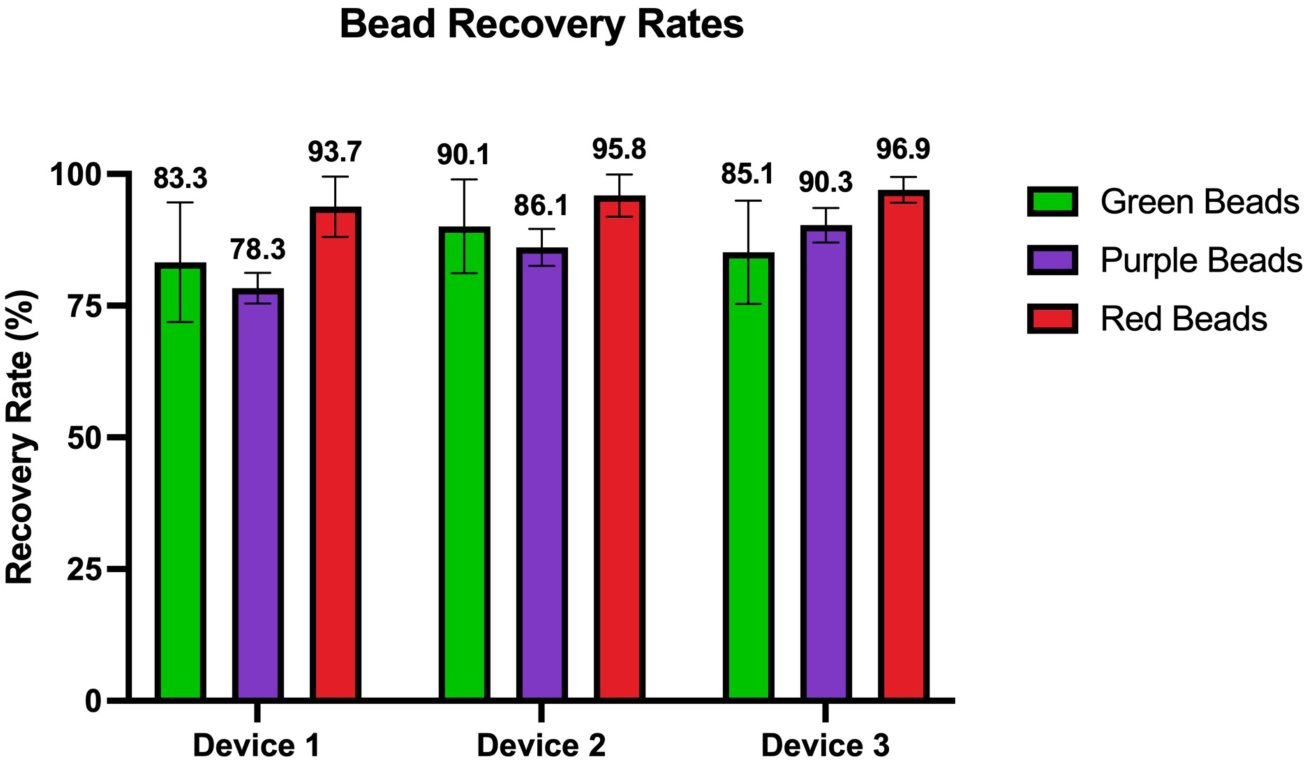
Recovery Rates across Different Magnetic Levitation Device Designs. The experimental recovery rate of green beads in the top outlet, purple beads in the middle outlet, and red beads in the bottom outlet across different devices during sorting trials (n=4, n=4, and n=8 for Device 1, Device 2, and Device 3 respectively).

**Supplementary Video 1.**
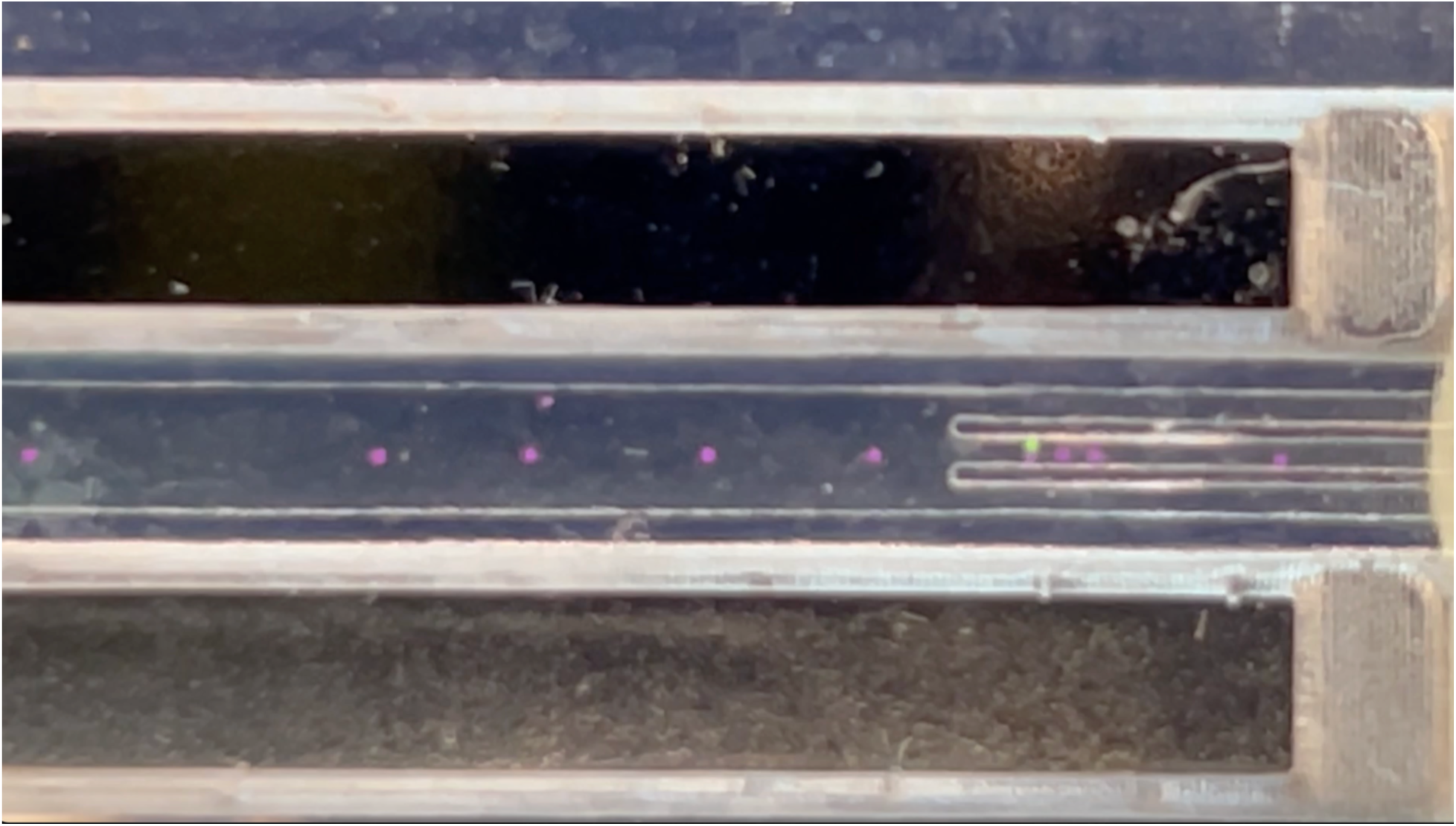
Polyethylene Bead Sorting. Sorting of 1.02, 1.06, and 1.09 g/mL polyethylene beads at 150 mM Gd in Device 3 with flow rates of 500 uL/min, 200 uL/min, and 400 uL/min across the top, middle, and bottom outlets respectively. Video taken on iPhone 14 Camera app used with 3x Zoom (4K resolution and 60 fps), and this clip was isolated from Device 3 Tracking Trial 5.

## References

1 P. H. Dannenberg, J. Kang, N. Martino, A. Kashiparekh, S. Forward, J. Wu, A. C. Liapis, J. Wang and S.-H. Yun, Lab. Chip, 2022, 22, 2343–2351.

2 K. D. Leake, B. S. Phillips, T. D. Yuzvinsky, A. R. Hawkins and H. Schmidt, Opt. Express, 2013, 21, 32605.

3 G. Zhu and N. Trung Nguyen, Micro Nanosyst., 2010, 2, 202–216.

4 D. Li, Encyclopedia of microfluidics and nanofluidics, Springer, New York, N.Y, 2008.

5 G. Choi, R. Nouri, L. Zarzar and W. Guan, Microsyst. Nanoeng., 2020, 6, 11.

6 H. Afsaneh and R. Mohammadi, Talanta Open, 2022, 5, 100092.

7 R. Salomon, S. Razavi Bazaz, W. Li, D. Gallego-Ortega, D. Jin and M. E. Warkiani, Micromachines, 2023, 14, 751.

8 H. Wei, B. Chueh, H. Wu, E. W. Hall, C. Li, R. Schirhagl, J.-M. Lin and R. N. Zare, Lab Chip, 2011, 11, 238–245.

9 P. Sajeesh and A. K. Sen, Microfluid. Nanofluidics, 2014, 17, 1–52.

10 Y. Song, R. Peng, J. Wang, X. Pan, Y. Sun and D. Li, ELECTROPHORESIS, 2013, 34, 684–690.

11 T. Kumar, A. V. Harish, S. Etcheverry, W. Margulis, F. Laurell and A. Russom, Lab. Chip, 2023, 23, 2286–2293.

12 S. Hettiarachchi, H. Cha, L. Ouyang, A. Mudugamuwa, H. An, G. Kijanka, N. Kashaninejad, N.-T. Nguyen and J. Zhang, Lab. Chip, 2023, 23, 982–1010.

13 Z. Zhao, S. Yang, L. Feng, L. Zhang, J. Wang, K. Chang and M. Chen, Adv. Mater. Technol., 2023, 8, 2202201.

14 A. J. Armstrong, M. S. Marengo, S. Oltean, G. Kemeny, R. L. Bitting, J. D. Turnbull, C. I. Herold, P. K. Marcom, D. J. George and M. A. Garcia-Blanco, Mol. Cancer Res., 2011, 9, 997–1007.

15 M. J. Tomlinson, S. Tomlinson, X. B. Yang and J. Kirkham, J. Tissue Eng., 2013, 4, 2041731412472690.

16 A. W. Wognum, A. C. Eaves and T. E. Thomas, Arch. Med. Res., 2003, 34, 461–475.

17 S. N. Jayasinghe, Adv. Biosyst., 2020, 4, 2000019.

18 S. Miltenyi, W. Müller, W. Weichel and A. Radbruch, Cytometry, 1990, 11, 231–238.

19 A. Dalili, E. Samiei and M. Hoorfar, The Analyst, 2019, 144, 87–113.

20 Y. Song, D. Li and X. Xuan, ELECTROPHORESIS, 2023, 44, 910–937.

21 C.-H. Hsu, D. Di Carlo, C. Chen, D. Irimia and M. Toner, Lab. Chip, 2008, 8, 2128.

22 F. Hossein and P. Angeli, Biophys. Rev., 2023, 15, 2005–2025.

23 A. H. Velders, L. Van Lieshout, E. A. T. Brienen, B. Diederich and V. Saggiomo, 2023, preprint, DOI: 10.26434/chemrxiv-2023-fnhkr-v2.

24 M. A. Witek, I. M. Freed and S. A. Soper, Anal. Chem., 2020, 92, 105–131.

25 H. Zhang and K.-K. Liu, J. R. Soc. Interface, 2008, 5, 671–690.

26 N. Abd Rahman, F. Ibrahim and B. Yafouz, Sensors, 2017, 17, 449.

27 K. Liang, S. Yaman, R. K. Patel, M. S. Parappilly, B. S. Walker, M. H. Wong and N. G. Durmus, 2022, preprint, DOI: 10.1101/2022.11.03.515127.

28 E. K. Chin, C. A. Grant, M. G. Ogut, B. Cai and N. G. Durmus, 2020, preprint, DOI: 10.1101/2020.07.27.223917.

29 N. Puluca, N. G. Durmus, S. Lee, N. Belbachir, F. X. Galdos, M. G. Ogut, R. Gupta, K. Hirano, M. Krane, R. Lange, J. C. Wu, S. M. Wu and U. Demirci, Adv. Biosyst., 2020, 4, 1900300.

30 N. G. Durmus, H. C. Tekin, S. Guven, K. Sridhar, A. Arslan Yildiz, G. Calibasi, I. Ghiran, R. W. Davis, L. M. Steinmetz and U. Demirci, Proc. Natl. Acad. Sci., DOI:10.1073/pnas.1509250112.

31 M. Anil-Inevi and E. Ozcivici, Springer US, New York, NY, 2025.

32 A. Nemiroski, A. A. Kumar, S. Soh, D. V. Harburg, H.-D. Yu and G. M. Whitesides, Anal. Chem., 2016, 88, 2666–2674.

33 X. Ren, M. C. Breadmore and F. Maya, Anal. Chem., 2024, 96, 3259–3266.

34 M. F. Alexandre-Franco, R. Kouider, R. Kassir Al-Karany, E. M. Cuerda-Correa and A. Al-Kassir, Micromachines, 2024, 15, 1137.

35 M. Levis, F. Ontiveros, J. Juan, A. Kavanagh and J. J. Zartman,.

36 K. Karimi, A. Fardoost, N. Mhatre, J. Rajan, D. Boisvert and M. Javanmard, Micromachines, 2024, 15, 1274.

37 L. C. Duarte, F. Figueredo, C. L. S. Chagas, E. Cortón and W. K. T. Coltro, Anal. Chim. Acta, 2024, 1299, 342429.

38 Hannah. B. Musgrove, Megan. A. Catterton and Rebecca. R. Pompano, Anal. Chim. Acta, 2022, 1209, 339842.

39 A. Bewley, Z. Ge, L. Ott, F. Ramos and B. Upcroft, in 2016 IEEE International Conference on Image Processing (ICIP), IEEE, Phoenix, AZ, USA, 2016, pp. 3464–3468.

